# SAMase of bacteriophage T3 inactivates *E. coli’s* methionine S-adenosyltransferase by forming hetero-polymers

**DOI:** 10.1101/2021.05.05.442785

**Authors:** Hadas Simon-Baram, Daniel Kleiner, Fannia Shmulevich, Raz Zarivach, Ran Zalk, Huayuan Tang, Feng Ding, Shimon Bershtein

## Abstract

S-adenosylmethionine lyase (SAMase) of bacteriophage T3 degrades the intracellular SAM pools of the host *E. coli* cells, thus inactivating a crucial metabolite involved in plethora of cellular functions, including DNA methylation. SAMase is the first viral protein expressed upon infection and its activity prevents methylation of the T3 genome. Maintenance of the phage genome in a fully unmethylated state has a profound effect on the infection strategy — it allows T3 to shift from a lytic infection under normal growth conditions to a transient lysogenic infection under glucose starvation. Using single-particle Cryo-EM and biochemical assays, we demonstrate that SAMase performs its function by not only degrading SAM, but also by interacting with and efficiently inhibiting the host’s methionine S-adenosyltransferase (MAT) — the enzyme that produces SAM. Specifically, SAMase triggers open-ended head-to-tail assembly of *E. coli* MAT into an unusual linear filamentous structure in which adjacent MAT tetramers are joined together by two SAMase dimers. Molecular dynamics simulations together with normal mode analyses suggest that the entrapment of MAT tetramers within filaments leads to an allosteric inhibition of MAT activity due to a shift to low-frequency high-amplitude active site-deforming modes. The amplification of uncorrelated motions between active site residues weakens MAT’s ability to withhold substrates, explaining the observed loss of function. We propose that the dual function of SAMase as an enzyme that degrades SAM and as an inhibitor of MAT activity has emerged to achieve an efficient depletion of the intracellular SAM pools.

**IMPORTANCE:** Self-assembly of enzymes into filamentous structures in response to specific metabolic cues has recently emerged as a widespread strategy of metabolic regulation. In many instances filamentation of metabolic enzymes occurs in response to starvation and leads to functional inactivation. Here, we report that bacteriophage T3 modulates the metabolism of the host *E. coli* cells by recruiting a similar strategy — silencing a central metabolic enzyme by subjecting it to phage-mediated polymerization. This observation points to an intriguing possibility that virus-induced polymerization of the host metabolic enzymes might be a common mechanism implemented by viruses to metabolically reprogram and subdue infected cells.

---

S-adenosylmethionine lyase (SAMase) is the first phage-induced protein to appear in the T3-infected cells (1). Although it has long been assumed that SAMase uses water molecules to break SAM into methylthioadenosine (MTA) and homoserine, a recent publication has shown that SAMase is, in fact, not a hydrolase but a lyase degrading SAM into MTA and homoserine lactone (2). Degradation of the intracellular SAM pools effectively subdues numerous SAM-utilizing reactions of the host cell, including RNA, DNA, protein and small molecule methylation, polyamine synthesis, and production of cofactors (3). Since SAM is an obligatory cofactor required for the action of restriction endonucleases belonging to type I R-M system (4), it was initially proposed that the primary biological role of SAMase expression was to render T3 bacteriophage immune against host restriction (5). However, this premise was later refuted by showing that phages lacking SAMase activity were nonetheless able to overcome host restriction systems just as efficiently as phages expressing the enzyme (6). Quite unexpectedly, SAMase was found to play a pivotal role in determining the infection strategy of T3 by maintaining its genome in a fully unmethylated state. While T3 is a virulent lytic phage when propagated in rich medium, it initiates a transient lysogenic infection in glucose-starved *E. coli* cells by blocking its own expression, provided that the infecting T3 DNA is unmethylated (7).

Given the central role played by SAMase in the infection strategy of T3 and the importance of SAM-utilizing reactions to cellular metabolism, it is unsurprising that multiple attempts have been made to purify and characterize this enzyme. It was reported early on that SAMase tightly binds to and co-purifies with a host protein factor, yet the identity of the protein or the significance of its interaction with SAMase remained obscure (8, 9).

## SAMase binds to and inhibits MAT

We began by identifying the unknown host factor that reportedly co-purifies with SAMase. To this end, we recombinantly expressed C-terminal His-tagged SAMase in *E. coli* and performed Ni-NTA affinity purification of the recombinant SAMase followed by size exclusion chromatography (see **Methods**). This purification procedure produced a mix of free dimeric SAMase molecules (~ 27 kDA) (**Fig. S1A-C**) and a high-molecular-weight complex comprised of SAMase molecules bound to an unknown host protein that, as previously reported(8, 9), appeared as a single band of ~43 kDA on SDS-PAGE (see **Fig. S1D**). By applying a shotgun MS analysis and an affinity pull-down assay, we determined that the *E. coli* protein that binds to and co-purifies with SAMase is methionine S-adenosyltransferase (MAT) (see **Methods, Table S1**, and **Fig. S1D,E**). *E. coli* MAT is a dihedral homotetramer (comprised of dimer of dimers) that catalyzes the condensation of ATP and methionine to produce SAM (10). MAT is an essential enzyme and the only source of SAM in *E. coli* (11). Thus, SAMase binds to the very enzyme whose unique product it degrades. To understand the functional significance of SAMase-MAT interaction, we measured how the interaction affected the activity of each of the enzymes *in vitro*. Whereas SAMase remained fully active in the presence of MAT, the activity of MAT was severely diminished and, ultimately, blocked with an increase in SAMase concentration (**Fig. 1**).

**Figure 1.**
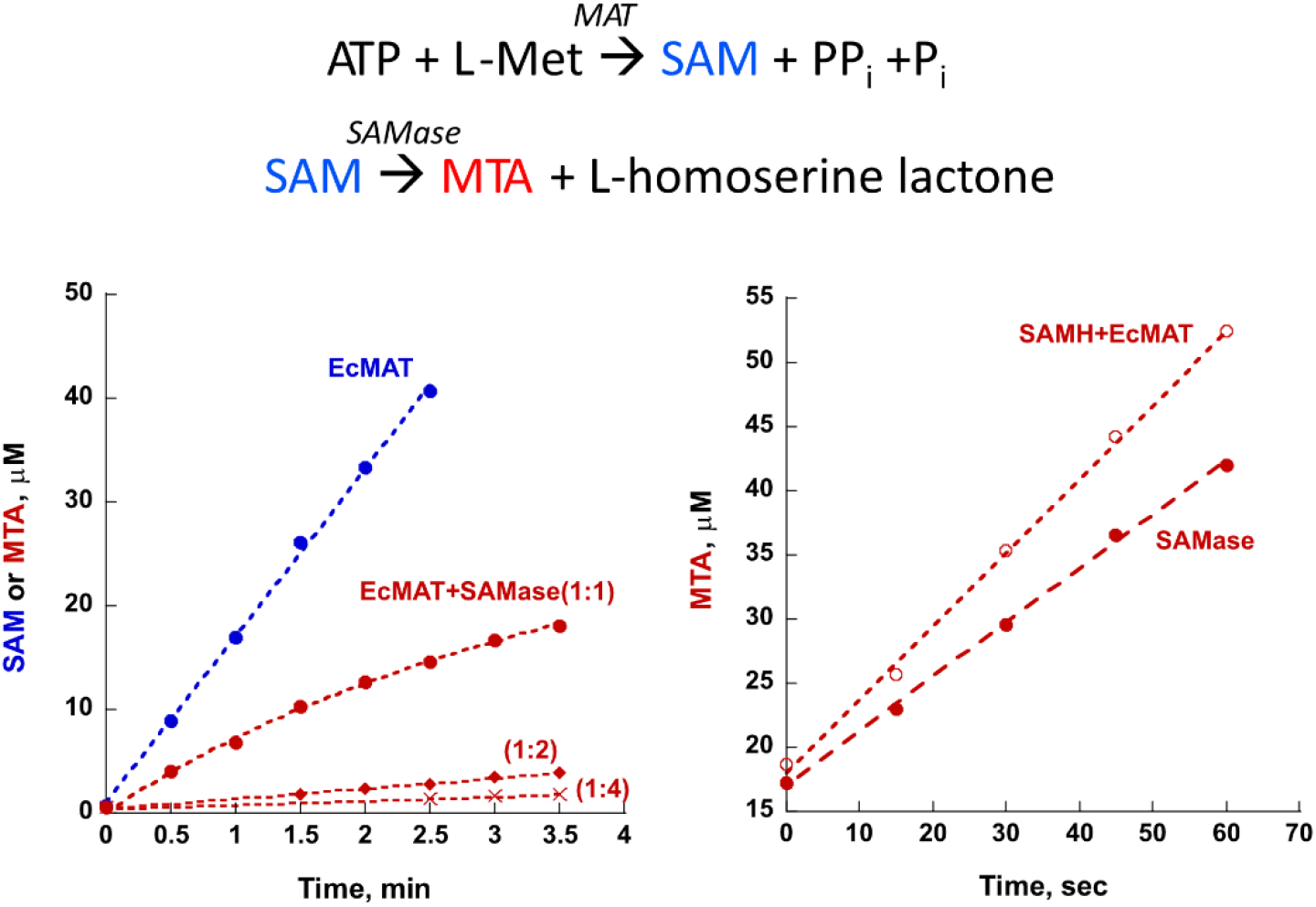
T3 SAMase inhibits *E. coli* MAT while preserving its own activity. *Left panel*. Synthesis of SAM from ATP and methionine by *E. coli* MAT was conducted in the absence (*blue* trace) and presence (*red* traces) of SAMase. The rate of SAM (*blue*) and/or methylthioadenosine (MTA) (*red*) production was monitored by HPLC (see **Methods** and **Fig. S11**). SAMase severely inhibits MAT activity already at the 1:1 EcMAT:SAMase molar ratio. Four-fold molar excess of SAMase almost fully diminishes the MAT activity. *Right panel.* SAMase activity was monitored by following MTA production upon addition of SAM in the absence (filled circles) or presence (empty circles) of *E. coli* MAT at 1:1 EcMAT:SAMase molar ratio. Note that no inhibition of SAMase activity is observed. The apparent improvement in the rate of SAM degradation in the presence of MAT might be attributed to the structural stabilization of SAMase.

## SAMase-MAT polymerization

To unravel the structural basis of *E. coli* MAT inactivation by T3 SAMase, we turned to single-particle Cryo-EM. To this end, recombinantly expressed and purified SAMase and MAT were mixed at 1:1 molar ratio in the presence of SAM and pre-incubated for 48 hours. The mix was then separated by size exclusion chromatography, and the high-molecular-weight peak comprised of both enzymes was subjected to Cryo-EM analysis (see **Methods** and **Fig. S2**). The collected images revealed filamentous structures of various lengths (**Fig. S3A**). Most 2D classes extracted from the images included both MAT and SAMase molecules (**Fig. S3B**). We solved the structure at 3.6Å resolution (PDB ID 70CK) (**Fig. S3C,D** and **Table S2**) and found that the filaments are formed of tetrameric MAT molecules brought together by SAMase dimers (**Fig. 2A**). Specifically, each SAMase monomer forms an interaction with a MAT monomer and another SAMase monomer, so that each two MAT tetramers are joined in a head-to-tail fashion by two SAMase dimers. The interaction between the adjacent MAT molecules is weak, with an interface of only ~170Å (see **Methods** and **Table S3**). A SAMase monomer is an α/β protein composed of nine anti-parallel β-strands forming a barrel-like structure and two helixes positioned outside the barrel (**Fig. 2B** and **Fig. S4A**). The topology of T3 SAMase fold appears to be unique, even in comparison to a recently solved phage-encoded Svi3-3 SAMase (2) (PDB ID 6zmg) (**Fig. S4B**). Two T3 SAMase monomers interact along ~1,000Å^2^isologous interface to form a C2-dimer. The MAT-SAMase interface is smaller and occupies ~600-700Å^2^(see **Methods** and **Table S3**). The “cracked egg” structure of SAMase dimer pushes one of the two joined MAT tetramers away from the plane (**Fig. 2B**). This bend between two adjacent MAT tetramers causes a propagation of right-handed helical twist along the entire SAMase-MAT heteropolymer. *In silico* assembly of the SAMase-MAT filamentous structure revealed that SAMase-MAT complex completes a full helical turn every nine MAT tetramers (**Fig. S5** and **Fig. 2C**). However, the distribution of MAT tetramers within the individual SAMase-MAT filaments found in Cryo-EM samples spanned only 2-7 units (**Fig. S3E**), which explains why we could not use the assumption of helical symmetry while solving the filamentous structure (see **Methods**). Since SAMase-mediated polymerization of MAT proceeds through an open-ended assembly, it is possible that the actual length of the filaments in solution (or *in vivo*) is, in fact, much longer than the distribution of lengths obtained in Cryo-EM analysis. Longer filaments probably form entangled structures of reduced solubility. Indeed, a mix of SAMase and MAT proteins subjected directly to Cryo-EM analysis without prior separation on a gel filtration column produced clumps of aggregates.

**Figure 2.**
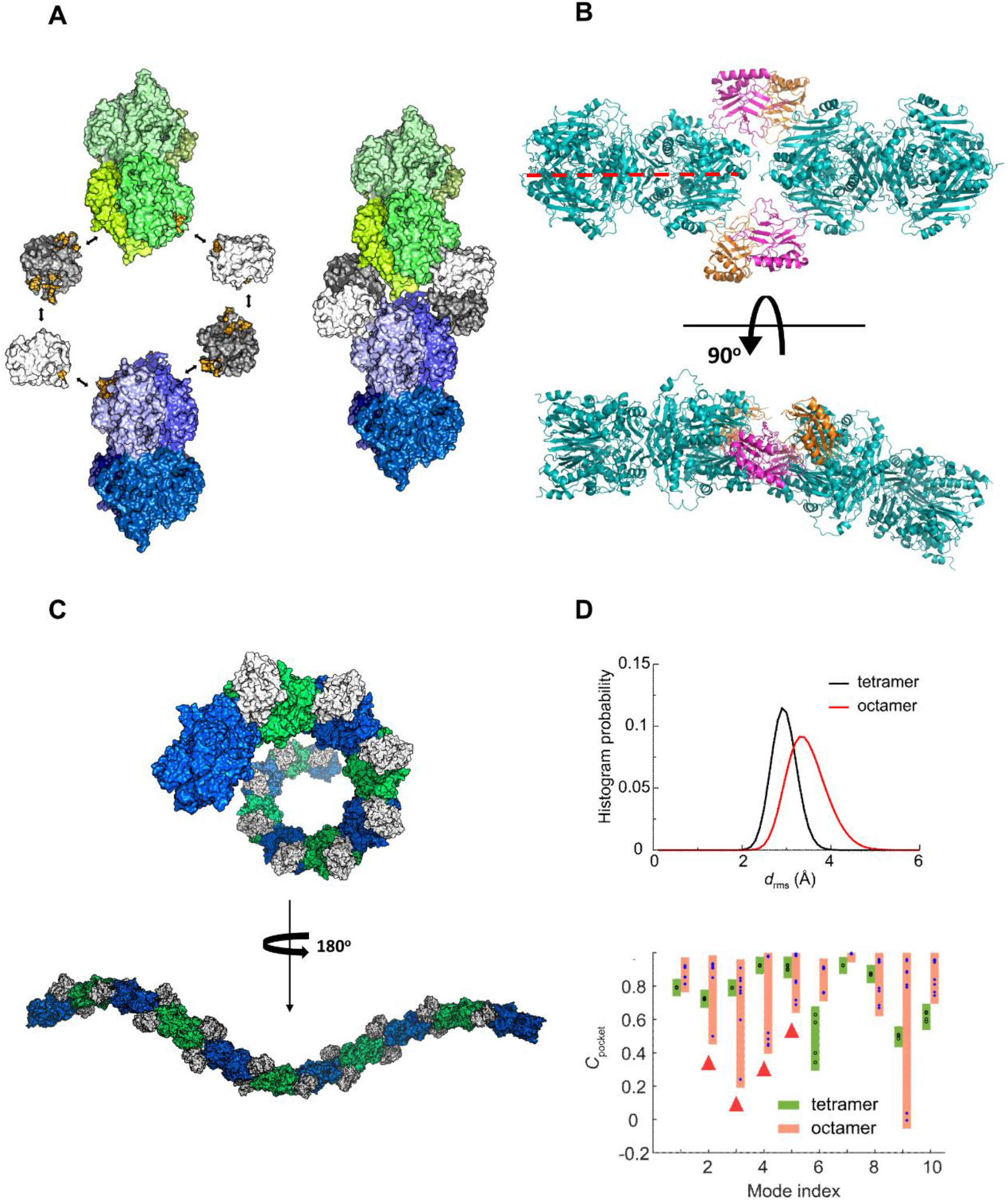
T3 SAMase triggers hetero-polymerization and inactivation of *E. coli* MAT. **A.** Two *E. coli* MAT tetramers (individual monomers are depicted in different shades of *green* and *blue*) are brought together by two SAMase dimers (shown in *white* and *grey*). The interfaces between the proteins are colored *yellow*. The properties of the interfaces are summarized in **Table S3.** Each SAMase monomer interacts with a MAT tetramer and another SAMase monomer. **B.** Rotating the SAMase-MAT hetero-oligomer 90° along the axis perpendicular to the dimeric interface of one of the MAT tetramers (*red* dashed line) reveals a ~30° kink in the assembly of two MAT monomers. MAT tetramers are shown in *cyan*. SAMase monomers are shown in *magenta* and *orange*. **C**. The *in silico* reconstructed SAMase-MAT filament reveals helical symmetry (the helix completes full turn every nine MAT tetramers). MAT tetramers are colored intermittently in *green* and *blue*. SAMase dimers are shown in *white*. See also **Fig. S5**. **D**. DMD simulations and normal mode analyses. *Upper panel*: Root-mean-square deviation *d_rms_* of tetramer (black line) and octamer (red line) pocket residues calculated from coarse-grained simulations. ‘Tetramer’ corresponds to active site residues in an unbound MAT tetramer. ‘Octamer’ corresponds to active site residues within two MAT tetramers bound by two SAMase dimers. *Lower panel:* Mean inter-residue dynamic correlation of pocket residues, *C_pocket_*, as a function of normal mode index. The lower *C_pocket_* value the more pocket deformation there is (when *C_pocket_* =1, there is no deformation). Four pockets in the tetramer are shown in circles and boxed in *green*, eight pockets in the octamer are shown in stars and boxed in *orange*. Pocket deformations in the octamer takes place at lower-index modes with significantly lower frequencies and higher amplitude (highlighted by *red* triangles) compared to the tetramer (see also **Fig. S9**).

## SAMase-MAT interaction is specific

To validate the functional relevance of the obtained structure, we identified twenty-one MAT residues participating in interface formation with SAMase monomers (see **Methods** and **Table S3, Fig. S6)**. Many of these residues appear to be conserved even among distantly related orthologous MATs whose activity we found to be unaffected by the presence of SAMase, such as MATs from *N. gonorrhea* or *U. urealyticum* (68% and 44% amino acid sequence identity with *E. coli* MAT, respectively) (**Fig. S6**, **Fig. S7A,B**). However, six out of the twenty-one interface-forming residues, Asp 132, Val 133, Ile 178, Ile 211, Ala 214, and Trp 26, are present only in *E. coli* or in other closely related enteric bacteria (**Fig. S6**). We reasoned that the replacement of these residues to their homologs from a SAMase-insensitive MAT should break the MAT-SAMase interaction and, thus, release the inhibition of *E. coli* MAT activity. To test this assumption, we generated an *E. coli* to *N. gonorrhea* MAT mutant carrying the following substitutions: Asp132Pro, Val133Thr, Ile178Val, Ile211Val, Ala214Pro, and Trp26Leu. As expected, these substitutions have fully removed the inhibitory effect of SAMase (**Fig. S7C**). Furthermore, dynamic light scattering (DLS) performed on *E. coli*, *N. gonorrhea*, and mutant *E. coli* MATs in the absence or presence of SAMase clearly indicated that the introduced substitutions eliminated the complex formation between *E. coli* MAT and SAMase (**Fig. S8**). Thus, SAMase-mediated filamentation of *E. coli* MAT is responsible for its functional inactivation.

## Inhibition of MAT activity is allosteric

To better understand the mechanism of MAT inhibition, we performed coarse-grained discrete molecular dynamics (DMD)(12) simulations of a MAT tetramer and a MAT octamer joined by two SAMase dimers (see **Methods**). We identified *i*) MAT pocket (active site) residues, using 6 Å expansion from the active site substrates (AMPPNP and methionine in MAT tetramer from PDB (ID: 1P7L)); and *ii*) residues forming the interface between MAT and SAMase, using distance cutoff of 0.85 nm (**Fig. S9A**). Comparison of root-mean-square deviations (*d*_rms_) of active site residues within an unbound MAT tetramer (’tetramer pockets’) and active sites residues within two MAT tetramers joined by two SAMase dimers (’octamer pockets’) calculated from coarse-grained simulations revealed that *d*_rms_ of octamer pocket residues were larger and broadly distributed than those of the tetramer (**Fig. 2D**), suggesting that the pocket was less stable and more deformed after filamentation. To determine the mechanism of pocket deformation in the octamer, we next performed normal mode analysis(13). Comparison of top normal modes suggested that the octamer featured lower-frequency and higher-amplitude motions than the tetramer (**Fig. S9B**). For each normal model, we defined the overall deformation of MAT pockets by mean inter-residue dynamic correlation of pocket residues (*C_pocket_*) and found that the deformation of pockets was significantly larger (low C*_pocket_* values) in the octamer in comparison to the tetramer at low-frequency and large-amplitude modes, which is consistent with our coarse-grained simulations (**Fig. 2D** and **Fig. S9B-C).** This extent of active site deformation potentially weakens MAT affinity to substrates and explains the observed loss of MAT function within filaments. In addition, we calculated the motion of MAT-SAMase interface residues in top normal modes (**Fig. S9A and D**). The deformation of the interface residues was described by mean inter-residue dynamic correlation, C*_interface_*. We found that the interfacial residues in the unbound tetramer undergo high-amplitude out-of-phase motion (low C*_interface_* values) at the low-frequency modes (**Fig. S9D**). However, these motions were severely inhibited in the octamer, suggesting that filamentation further amplifies pocket deformation(14). Collectively, DMD simulations together with normal mode analyses suggest that the entrapment of MAT tetramers within the SAMase-mediated filaments leads to an allosteric inhibition of MAT activity.

## Conclusions

Maintenance of the T3 genome in a fully unmethylated state is the single determining factor in the phage ability to establish a transient lysogenic infection upon infection of glucose-starved *E. coli* cells (7). Considering the importance of the methylation status of the T3 phage genome to its infection strategy, the dual function of SAMase as a SAM-degrading enzyme *and* as an inhibitor of the only enzyme that produces SAM in the host cell provides a clear evolutionary advantage, as it ensures an efficient prevention of phage genome methylation. To our knowledge, the mechanism by which T3 SAMase inhibits the activity of *E. coli* MAT, *i.e.*, by mediating the polymerization of MAT tetramers, was not previously reported to any viral protein. Strikingly, however, it resembles the wide-spread phenomenon of filamentation of central metabolic enzymes in organisms as distinct as bacteria, yeast, worm, drosophila, and human (15). In approximately half of the cases in which the biological role of enzyme filamentation was determined, it was found that filamentation inhibits the enzyme activity, particularly when filamentation was triggered by starvation. In other cases, filamentation led to enzyme activation or even a change in enzyme specificity (15). As our understanding of the regulation of cellular metabolism by means of enzyme filamentation continues to grow, it becomes apparent that the advantage of metabolic filamentation is rooted in its ability to rapidly affect metabolic function in response to environmental transitions without changing enzyme concentration. Our finding that a viral protein can rapidly exert a metabolic control over the host cell by recruiting a similar strategy of metabolic filamentation opens an interesting possibility that filamentation of host enzymes might be a wide-spread strategy implemented by viruses to reprogram and subdue the metabolic state of the host.

## Methods

### SAMase cloning and expression

A T3 SAMase encoding gene (UniProt P07693) with the addition of six His-encoding codons at 3’ was synthesized *de novo* by Integrated DNA Technologies (IDT) and cloned into pZA31 plasmid (Expressys) by Gibson assembly, under P_LTetO-1_ promoter (16). The resulting vector was transformed into a DH5αZ1 *E. coli* strain, which constitutively expresses Tet repressor, thus tightly quenching leaky expression (16). To express the protein, an overnight starter of the strain grown in LB media at 37°C in the presence of 30 μg/ml chloramphenicol was diluted 1:100 into YT media supplemented with 30 μg/ml chloramphenicol and grown at 37°C to O.D. at 600nm of ~0.6. At this point, SAMase expression was induced by adding 0.2 μg/mL anhydrotetracycline (aTc). Upon addition of the inducer, the temperature was reduced to 20°C, and the cells were allowed to grow overnight. The cells were then harvested by centrifugation (5,000 g, 20 min) and subject to either native or denatured purification protocols.

### SAMase purification under native conditions

For purification under native conditions, the harvested cells were pre-incubated for 30 min on ice with 1 mg/ml lysozyme, sonicated, centrifuged, and the resulting soluble fraction was applied on a Ni-NTA column (GE Healthcare, His Trap FF 1 ml) and purified according to the manufacturer’s instructions. The eluted protein was dialyzed at 4°C against Buffer A (20 % glycerol, 20 mM Hepes pH7, 150 mM KCl, and 5 mM DTT), concentrated (Amicon centrifugal filter Units, 3kD cut-off), and applied on a gel filtration column (Superdex 200 Increase 10/300 GL column, GE healthcare) pre-equilibrated with Buffer B (25 mM Tris pH 8.0, 150 mM KCl, 1 mM DTT). SAMase eluted at 18.4 ml, which corresponded to ~24 kDa (see **Fig. S1A,B**). The purified SAMase was dialyzed against Buffer A and stored at −20°C in Buffer A brought to 50% glycerol.

### SAMase purification under denatured conditions

Only a small fraction of SAMase purified under native conditions was found in a free unbound form (see **Fig. S1A**). To generate sufficient amounts of free SAMase, we performed purification under denatured conditions followed by refolding. To this end, the harvested cells were lysed in chilled lysis buffer (6 M Guanidine Hydrochloride, 20 mM Sodium Phosphate, pH7.8, 500 mM NaCl) for 1 hour and centrifuged at 5,000 g. The lysate supernatant was than loaded on a Ni-NTA column (GE Healthcare, His Trap FF 1 ml) pre-equilibrated with a binding buffer (8 M Urea, 20 mM Sodium Phosphate pH7.8, 500 mM NaCl). The column was washed with 10 column volumes of the binding buffer. SAMase was eluted with an elution buffer (8 M Urea, 20 mM Sodium Phosphate pH4, 500 mM NaCl). The eluted protein sample was diluted to > 0.05 mg/ml and dialyzed once against 6M urea, 20 mM Hepes pH7,150 mM KCl, 5 mM DTT and 20% glycerol at 4°C. The sample was then dialyzed three more times against buffer A at 4°C. Upon completion of dialysis, the sample was centrifuge at 12,000 g for 30 min. The supernatant was collected and concentrated using a filtration unit (Amicon centrifugal filter Units, 3kD cut-off). The purified refolded SAMase was stored at −20°C in Buffer A brought to 50% glycerol. The refolded SAMase was functional (*k_cat_* = 87.4 sec^−1^, K_m_ = 6 M) and produced an elution pattern on a gel filtration column similar to that of the natively purified protein (elution volume 18.1 ml) (**Fig. S10**).

### Determining the oligomeric status of SAMase in solution

Based on the primary sequence, the anticipated size of SAMase monomer is around 17kDA. The estimation of SAMase size by a gel filtration column using beta amylase (200 kDA), alcohol dehydrogenase (150 kDA), albumin (66 kDA), carbonic anhydrase (29 kDa), and cytochrome C (12.4 kDa) as protein standards produced a molecular mass of ~24-27 kDA for both natively purified and refolded SAMase (see **Fig. S1B** and **S10A**), suggesting that SAMase may form a dimer in solution. To test this possibility, we performed a cross-linking analysis. Specifically, 28 μM of SAMase were mixed with glutaraldehyde (Sigma, #G6257) to a final concentration of 2.5, 5 or 10% v/v. The protein mixes were then incubated for 30 min at 25 °C. Quenching was performed by the addition of one sample volume of 1M Tris-HCl at pH 8.0 and half sample volume of X5 concentrated Laemmli sample buffer. Finally, the quenched samples were boiled for 5 min at 60°C and separated by SDS-PAGE. Bands corresponding to dimeric (and, to a lesser extent, tetrameric) species appeared in all chosen glutaraldehyde concentration, suggesting that SAMase exists primarily as a dimer in solution (**Fig. S1C**).

### Activity assays

All enzymatic assays were performed at 37°C in activity buffer (25mM Hepes pH7.5, 100mM KCl, 10mM MgCl_2_, 1mM DTT). Activity of MATs in the absence of SAMase was determined at a saturated concentration of methionine (1 mM), 80μM ATP and 250nM enzyme. The enzymatic reaction was initiated by adding ATP. To measure MAT activity in the presence of SAMase, MATs were pre-incubated with SAMase at 1:1 to 1:4 molar ratios (250nM MAT and 250nM – 1μM SAMase) for 30 min prior to addition of ATP. To measure the effect of *E. coli* MAT on SAMase activity, 2μM SAMase were pre-incubated for 48 hours at 4°C either alone or in the presence of 2μM of *E. coli* MAT. The pre-incubated mixes were then diluted 1/40 in the activity buffer and SAMase activity was measured by adding 80μM SAM. SAMase catalytic parameters in the absence of MAT were determined at a range of SAM concentrations (0-150μM). SAMase (1μM) was pre-incubated for 30 min with 2μM BSA (as a crowder) prior to addition of SAM. Aliquots were removed from the reaction mixes at various time points (up to 4.5 min, every 30 sec), and the reactions were stopped by mixing with 10% perchloric acid in a 1:1 ratio. The aliquots were then centrifuged, and the supernatant was separated by high-performance liquid chromatography (HPLC) using the MultiHigh SCX 5μ 250 × 4.6 mm column (CS-Chromatographie Service GmbH). The mobile phase consisted of 400 mM ammonium fumarate (adjusted to pH 4.0 using formic acid) at a flow rate of 1 ml/min, while measuring absorbance at 254 nm. Data analysis was performed by integrating the peaks corresponding to SAM and methylthioadenosine (MTA) and fitting them to the corresponding calibration curves (**Fig. S11**). The kinetic constants were derived by fitting the resulted data point to a Michaelis-Menten equation (Vmax·[S]/Km+[S]).

### MS

Protein bands of ~43 kDa were incised from Coomassie stained SDS-PAGE gel (**Fig. S1D**) and digested in-gel by trypsin according to the manufacturer’s protocol (Promega). Peptides were then extracted and subjected to LC/MS analysis using an Eksigent nano-HPLC connected to the LTQ Orbitrap XL (Thermo Fisher Scientific). Reverse-phase chromatography of peptides was performed using an Acclaim PepMap Thermo Fisher scientific (C 18-column, 15 cm long, 75 μm ID, packed with 300 Å, 5 μm beads). Peptides were separated by a 70-min linear gradient, starting with 100% buffer A (5% acetonitrile, 0.1% formic acid) and ending with 80% buffer B (80% acetonitrile, 0.1% formic acid), at a flow rate of 300 nl/min. A full scan, acquired at 60,000 resolution, was followed by CID MS/MS analysis performed for the five most abundant peaks, in the data-dependent mode. Fragmentation (with minimum signal trigger threshold set at 1,000) and detection of fragments were carried out in the linear ion trap. Maximum ion fill time settings were 300 ms for the high-resolution full scan in the Orbitrap analyzer and 100 ms for MS/MS analysis in the ion trap. *E. coli* MAT protein was identified and validated using the SEQUEST search algorithms against *E. coli* (NCBI proteome database collection) operated under the Proteome Discoverer 2.4 software (Thermo Fisher Scientific). Mass tolerance for precursors and fragmentations was set to 10 ppm and 0.8 Da, respectively.

### Affinity pool-down

*E. coli* cells were grown in 25 ml LB to OD at 600nm = 0.5, harvested by centrifugation (4,500*g* for 15 min at 4°C), and lysed with Bug Buster (Novagene # 70921) in 50mM NaH_2_PO_4_ pH8, 300mM NaCl, and 250 units of Benzonase^®^ Nuclease (Millipore) for 30 min at room temperature. The cell lysate was then centrifuged (12,000*g* for 10 min at 4°C), and the collected supernatant was mixed with 4μM of 6xHis-tagged SAMase. The mix was pre-incubated for 30 min at room temperature while rotating at 60 rpm. The mix was then applied on Ni-NTA Beads spin columns (G BioSciences,0.2 ml #786-943), pre-equilibrated with Binding buffer (50mM NaH_2_PO_4_ pH8, 300mM NaCl, and 10 mM imidazole). The column was then washed ×5 with 2 CV of Wash buffer (50mM NaH_2_PO_4_ pH8, 300mM NaCl, and 20 mM imidazole), and the protein sample was eluted with 3 CV of Elution buffer (50mM NaH_2_PO_4_ pH8, 300mM NaCl, and 10 mM imidazole). The eluted sample was separated on SDS-PAGE and analyzed by Western Blot using custom-raised rabbit anti-*E.Coli* MAT polyclonal antibodies (Genemed synthesis Inc) and goat anti-rabbit HRP secondary antibodies (Abm #SH026). HRP on the immunoblot was detected with ECL (Advansta) (see **Fig. S1E**)

### Preparation of protein samples for Cryo-EM analysis

Refolded SAMase (150μl of 160μM of monomer in buffer A) was mixed with 150μl of 160μM (of monomer) of *E. coli* MAT in 25mM Tris pH8.0, 1mM DTT, and 10% glycerol. The two proteins were gently mixed and supplemented with 500μM of SAM. The sample was incubated at room temperature for 1 hour and then stored for 48 hours at 4°C. Next, additional 500μM of SAM were added, and the sample was centrifuged at 12,000*g* for 20 min in 4 °C. The supernatant was then loaded on SEC column (Superdex 200 Increase 10/300 GL column, GE healthcare) pre-equilibrated with Buffer B. High-molecular-weight peak eluted between the void volume of the SEC column (~8 ml) and 11 ml and comprised of both MAT and SAMase was collected (see **Fig. S2**). The collected protein sample was then concentrated using a filtration unit (Amicon ultra centrifugal filters, 3kD cut-off) to reach a total protein concentration of 0.2 mg/ml. Cryo-EM samples were prepared by plunge-freezing into liquid ethane. 3μl of protein solution at a concentration of 0.2 mg/ml were deposited on glow-discharged Quantifoil R 1.2/1.3 holey carbon grids (Quantifoil Micro Tools GmbH, Germany). Samples were manually blotted for four seconds at room temperature and vitrified by rapidly plunging into liquid ethane using a home-built plunging apparatus. The frozen samples were stored in liquid nitrogen until imaging.

### Cryo-EM data acquisition, data refinement, and model building

Samples were loaded under cryogenic conditions and imaged in low dose mode on a FEI Tecnai F30 Polara microscope (FEI, Eindhoven) operated at 300 kV. Datasets were collected using SerialEM(17), using a homemade semi-automated data collection script (https://doi.org/10.1021/jacs.0c07565). Images were collected by a K2 Summit direct electron detector fitted behind an energy filter (Gatan Quantum GIF) set to ±10 eV around zero-loss peak. Calibrated pixel size at the sample plane was 1.1 Å. The detector was operated in a dose fractionated counting mode, at a dose rate of ~8 ē/pixel/second. Each dose-fractionated movie had 50 frames, with total electron dose of 80 ē/Å^2^. Data was collected at a defocus range of −1.0 to −2.0 μm. Dose-fractionated image stacks were aligned using MotionCorr2 (18), and their defocus values were estimated by Gctf(19). The aligned sum images were used for further processing by cryosparc v2.15.0 (https://doi.org/10.1038/nmeth.4169). Particles were “blob” picked followed by 2D classification. Auto-picked particles based on a template produced from a small subset of manually picked 2D class averages. The picked particles were processed by *ab initio* 2D classification followed by manual inspection and selection of the class averages. Classes were selected based on shape and number of particles. All selected classes had clear signatures of secondary structure elements. Initial 3D reference was prepared from the dataset of the “untreated” sample. Final maps were obtained by 3D refinement with no symmetry imposed (C1). Resolution was assessed by Fourier-Shell-Correlation (**Fig. S3**). The final cryo-EM maps following density modification were used for model building. The model of the MAT complex structure (PDB ID 3P7L) was used as an initial template and rigid-body-fitted into the cryo-EM density for the highest-resolution state of DNA-PKcs in UCSF Chimera (20) and manually adjusted and rebuilt in Coot (21). Real-space refinement were then performed in Phenix (22). SAMase structure was built *de-novo* into the cryo-EM density map. Following refinement of the highest-resolution state, The MAT-SAMase structure was deposited into the PDB with the codes shown in **Table S2**.

### Protein interface analysis

MAT-SAMase and SAMase-SAMase interfaces were characterized using protein interfaces, surfaces, and assembly (PISA) server (23). The interface area is calculated by PISA as a difference in total accessible surface areas of isolated and interfacing structures divided by two. Change in the solvation free energy upon formation of the interface, **Δ^i^G** in kcal/mol, is calculated as a difference in total solvation energies of isolated and interfacing structures. Negative **Δ^i^G** corresponds to hydrophobic interfaces and does not include the effect of satisfied hydrogen bonds and salt bridges across the interface. PISA also predicts the formation of hydrogen-bonds and salt-bridges in the interface. The individual parameters of each interface are summarized in **Table S3**.

### Dynamic Light Scattering (DLS)

MATs and SAMase were analyzed either separately (18.4μM MAT, 22μM SAMase) or together (2.85μM MAT and 11.4uM SAMase, 1:4 ratio). All measurements were done in buffer Tris pH8.0, 150mM KCl, and 1mM DTT. All samples were supplemented with 500μM SAM and pre-incubated for 30 min at room temperature prior to analysis. Spectra were collected in 1 ml volume by using CGS-3, (ALV, Langen, Germany). The laser power was 20mW at the He-Ne laser line (632.8nm). Averaged (10 runs) scattered intensities were measured by ALV/LSE 5004 multiple tau digital cross correlator, at 60-90°, during 30s at room temperatures. The correlograms were fitted with version of the program CONTIN(24).

### Coarse-grained discrete molecular dynamics simulations

Coarse-grained simulations were performed with discrete molecular dynamics (DMD) simulations. The details of DMD algorithm can be found in previous publications (25). To reduce the computational cost, each residue was represented by one Cα bead. Residue pairs forming native contact (cutoff=7.5 Å) experienced attractive Gō potentials and hardcore repulsions were applied to residue pairs without native contacts (26). For each system, 39 independent simulations starting with different initial positions and velocities were performed. Each trajectory lasted 6 million DMD steps with only the last 4 million simulation data was used for equilibrium conformational analysis. The deformation of pocket was characterized by root-mean-square deviation 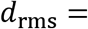 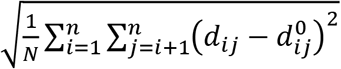, where *n* was the number of pocket residues, 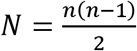 was the number of pocket residue pairs, 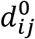 and *d*_*ij*_ were the distances between residue *i* and residue *j* in the native state and simulations, respectively.

### Normal model analyses

Normal mode analyses were conducted with oGNM server (13). Each residue was represented by one network node with native contacts connected by elastic springs. The residue motions between residue *i* and residue *j* were described by the correlation matrix 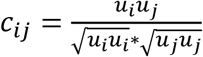, where *u* was the eigenvector of each mode. The amplitude of each mode was 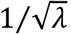, where λ was the eigenvalue of each mode. The overall deformations of pocket and interface residues were characterized by mean inter-residue dynamic correlation of residues, i.e., C*_pocket_* and *C_interface_*.

## Acknowledgements

We acknowledge funding to S.B. from Israel Science Foundation (personal grant 1630/15) and to F.D. from NIH (R35GM119691).

---

**Table S1:** Mass spectrometry analysis.

|  |  |
| --- | --- |
| Accession | QRB62406.1 |
| Description | methionine adenosyltransferase<br>[Escherichia coli] |
| Coverage [%] | 71 |
| # Peptides | 24 |
| # PSMs | 240 |
| # Unique Peptides | 24 |
| # AAs | 384 |
| MW [kDa] | 41.9 |
| calc. pI | 5.26 |
| Score Sequest HT: Sequest HT | 787.55 |
| # Peptides (by Search Engine):<br>Sequest HT | 24 |
| # Protein Pathway Groups | 0 |
| Found in Sample: [S1] F1: Sample | High |
| Found in Sample: [S2] F2: Sample | High |
| # Protein Groups | 1 |

**Table S2:**
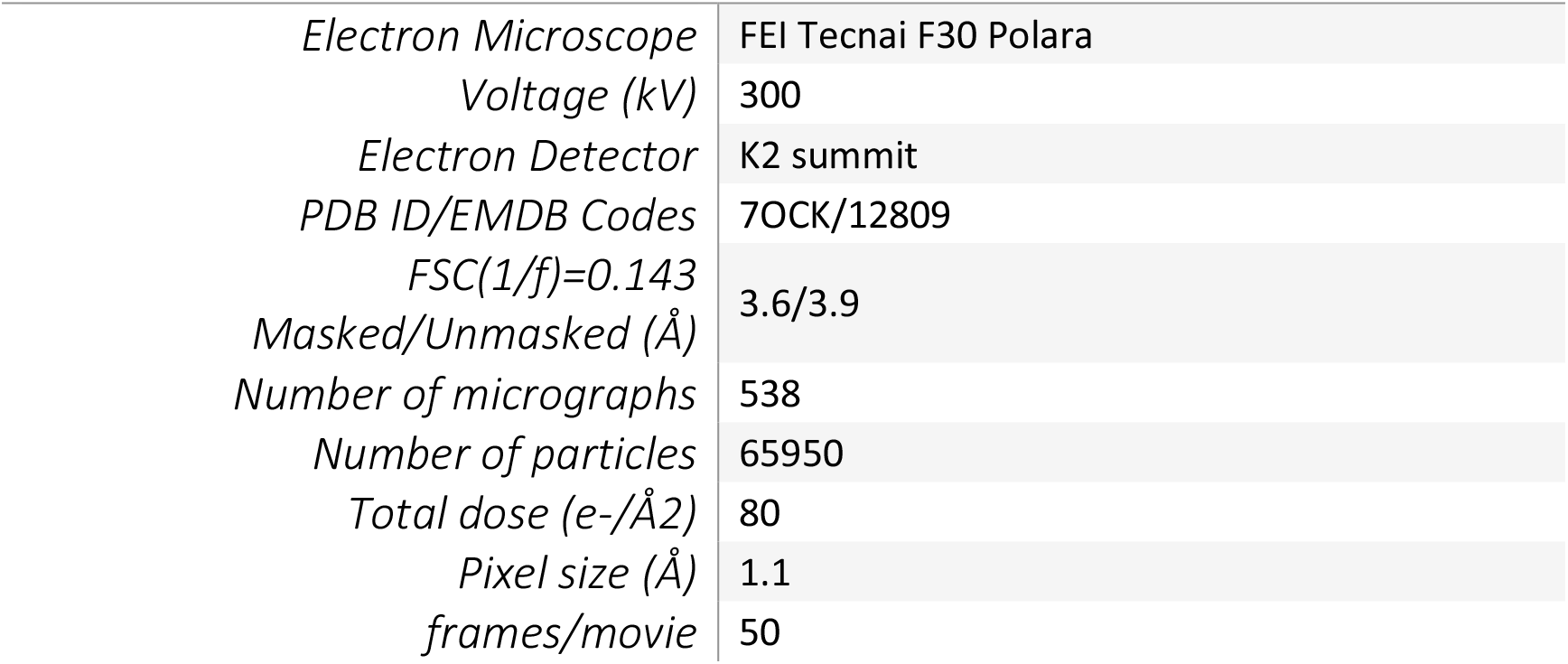
Data collection and refinement parameters of the cryo-EM reconstruction in this study.

**Figure S1.**
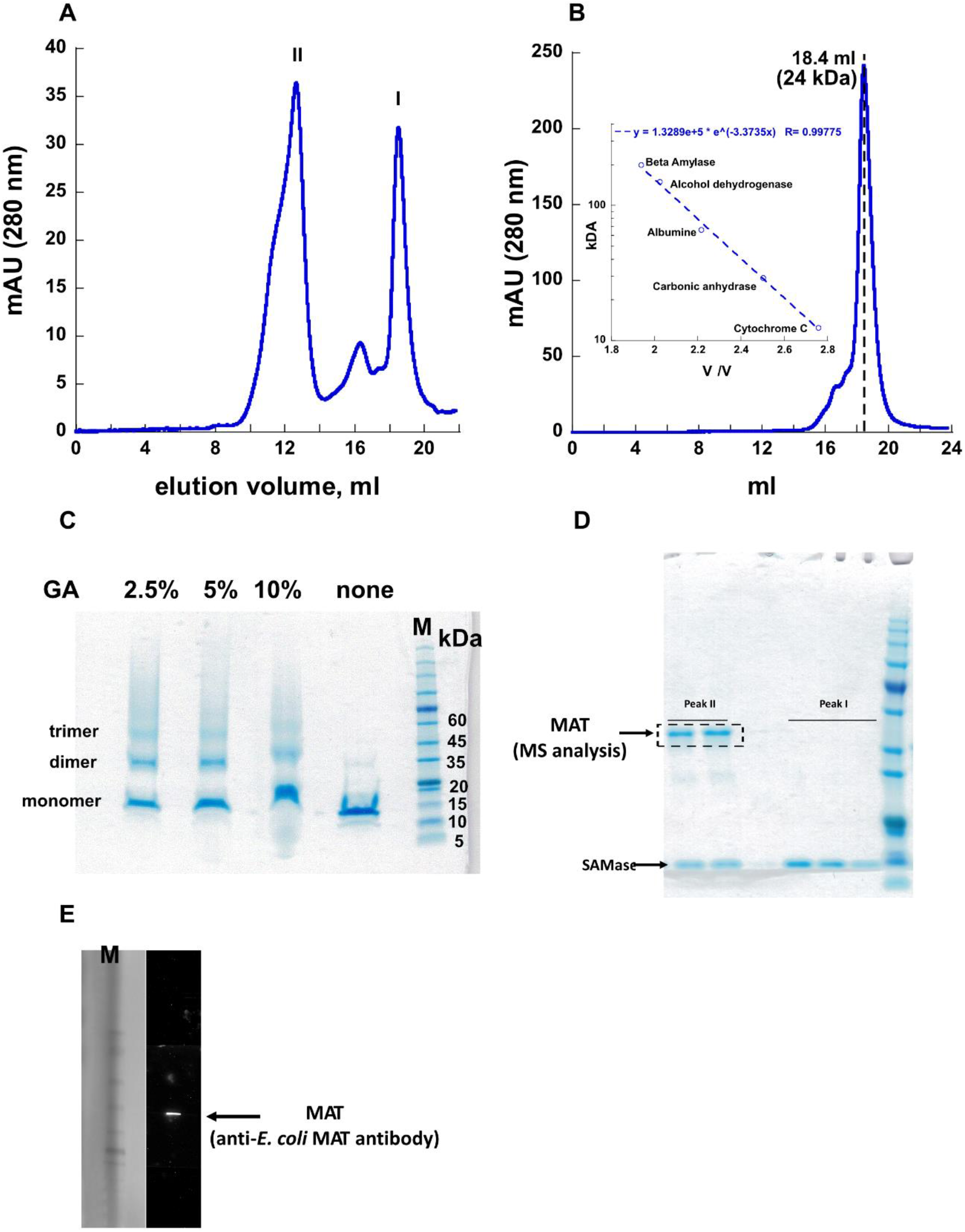
Purification of SAMase under native conditions. **A.** Size exclusion chromatography following Ni-NTA purification of recombinantly expressed 6xHis-tagged SAMase under native conditions (see **Methods** for details). Two major peaks are marked I and II. **B.** Free SAMase eluted as peak I was concentrated and re-run on the same SEC column. The protein eluted as a single peak at 18.4 ml that, according to molecular weight protein markers (inset), corresponds to 24 kDa. **C.** SDS-PAGE analysis of SAMase cross-linked with 2.5%, 5%, or 10% v/v glutaraldehyde (GA). Dimeric species are easily distinguished, suggesting that SAMase exists as a dimer in solution (see **Methods** for details). **D.** Proteins eluted with peaks I and II were collected at several elution aliquots and analyzed in SDS-PAGE. SAMase was found in both peaks. A higher-molecular-weight protein (around~43 kDa) was found only in peak II. The protein was excised from the gel (dashed lines) and subjected to MS analysis (see **Methods**). **E**. Western-Blot detection with custom raised polyclonal antibodies of *E. coli* MAT pulled-down from a cell lysate with SAMase (see **Methods**). Marker is shown on the left.

**Figure S2.**
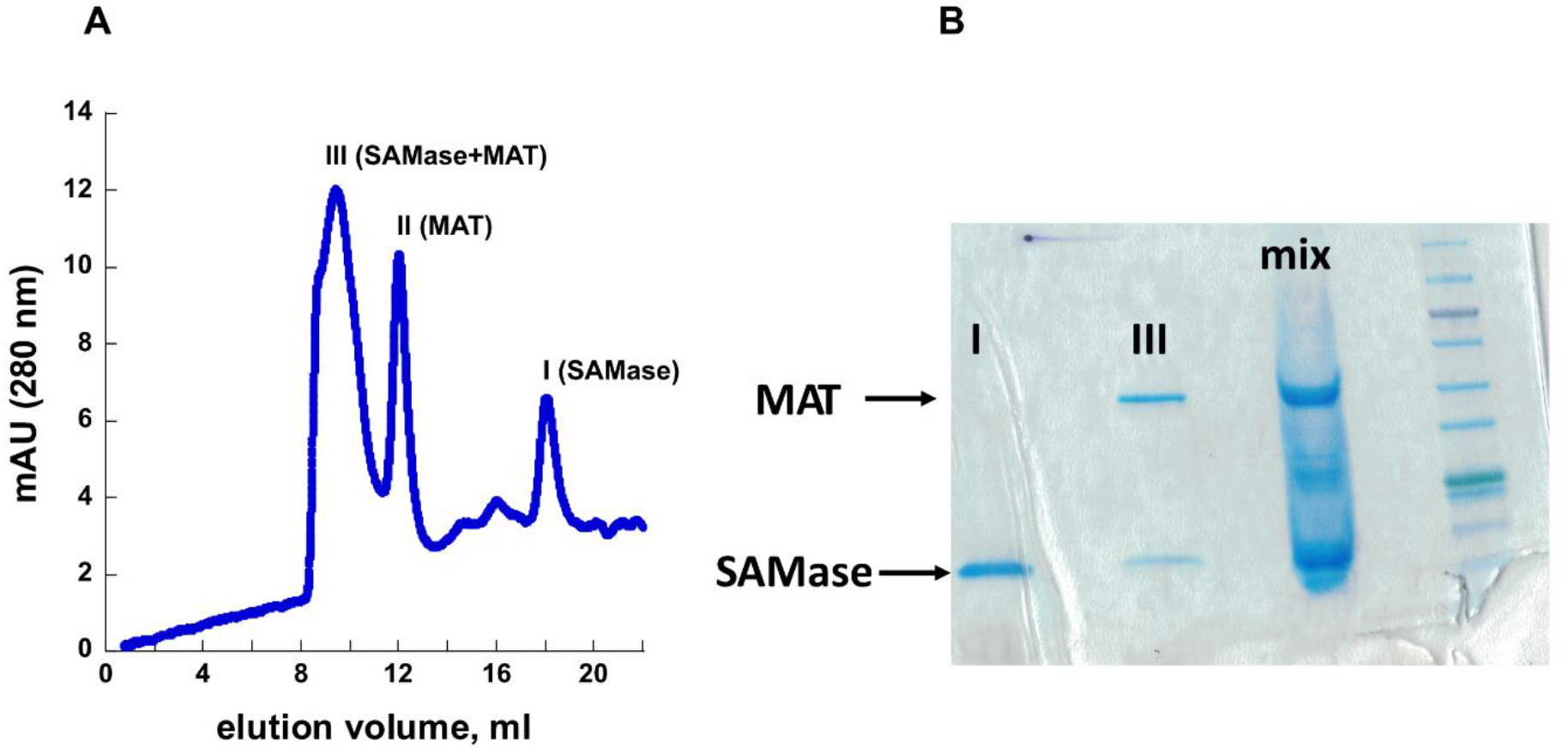
Preparation of SAMase-MAT mix for Cryo-EM analysis. **A.** Size Exclusion Chromatography of refolded SAMase and *E. coli* MAT mix pre-incubated in the presence of SAM (see **Methods** for details). Three major peaks are marked: I (SAMase only), II (MAT only), and III (SAMase+MAT). **B.** SDS-PAGE analysis of SAMase-MAT mix. “Mix” corresponds to a mix of SAMase and MAT prior to Size Exclusion Chromatography. The protein composition of eluted peaks: Peak I (SAMase only), Peak III (both SAMase and MAT). Marker is shown on the right.

**Figure S3.**
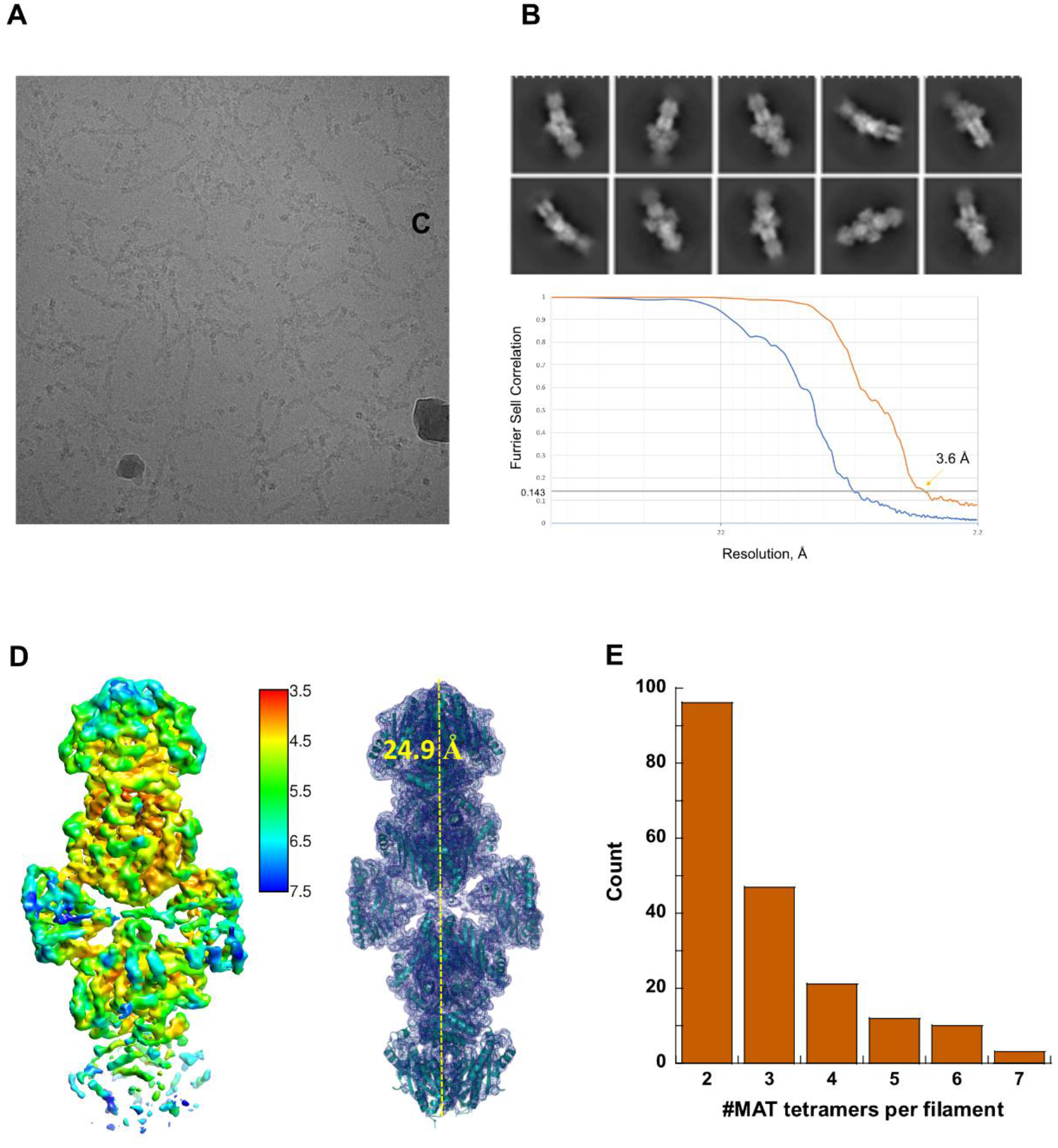
Cryo-EM reconstruction of T3 SAMase – *E. coli* MAT complex. **A.** A representative Cryo-EM micrograph of SAMase-MAT filaments. **B.** 2D class averages. **C.** The resolution of the structure is 3.6Å by gold standard FSC. **D.**, *left panel*: Local resolution map of two MAT tetramers connected by two SAMase dimers; *right panel*: Ribbon representation of the structure on the left embedded in the electron density map. The distance between the two most remote points of the opposing eMAT tetramers amounts to 24.9Å. **E.** Distribution of the number of MAT tetramers found in the individual SAMase-MAT filaments.

**Figure S4.**
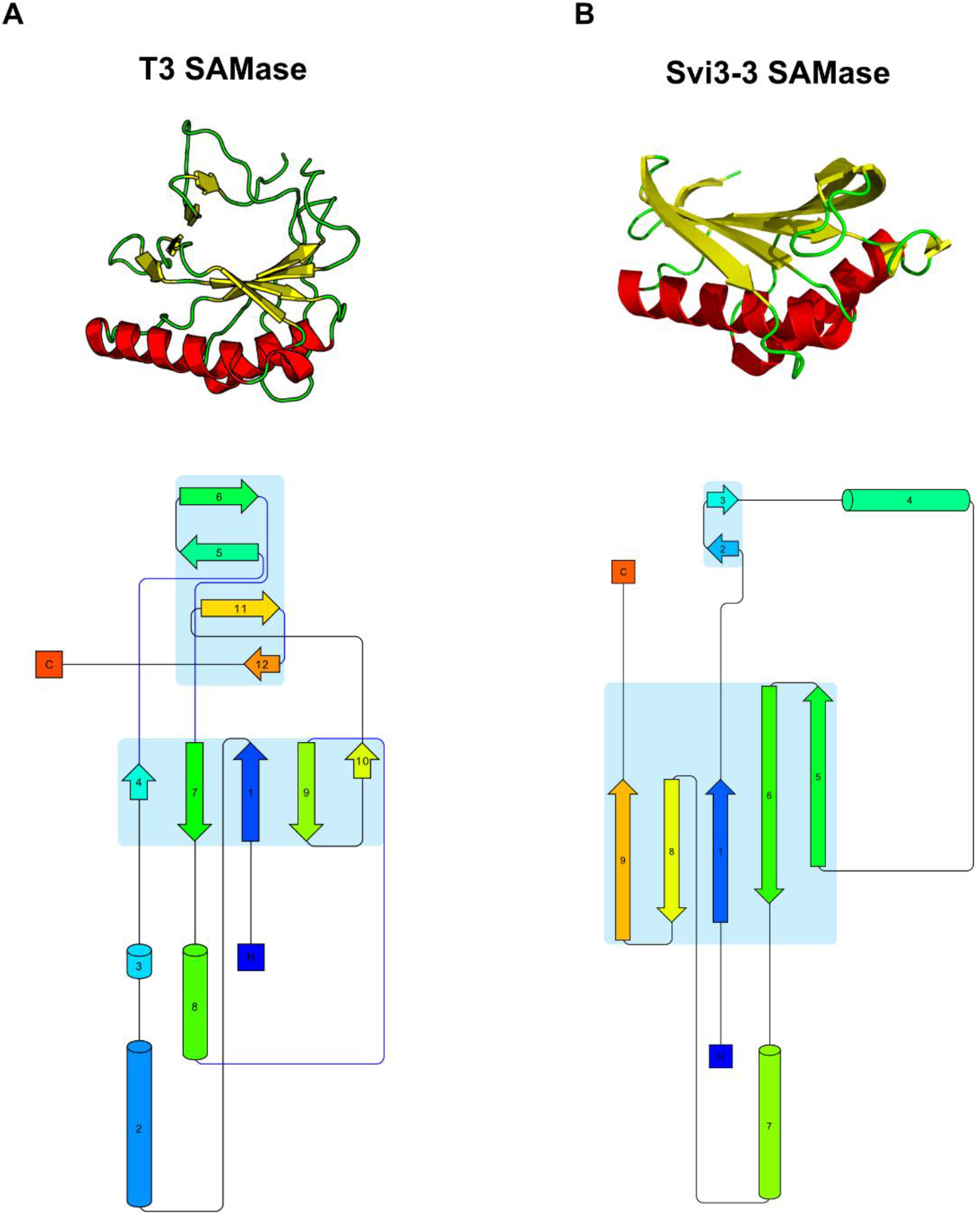
Comparison fold topologies of T3 SAMase and Svi3-3. **A.** *Upper panel:* Ribbon presentation of T3 SAMase monomer. Nine antiparallel β-strands (*yellow*) assemble into a barrel-like structure and two α-helixes (*red*) are placed outside the barrel. *Lower panel:* Secondary structure elements connected along the primary sequence reveal fold topology of T3 SAMase monomer. Arrows and cylinders represent betta-strands and alpha-helixes, respectively. The blue shading depicts betta-pleated sheets**. B.** *Upper panel:* Ribbon presentation of Siv3-3 SAMase monomer (PDB ID: 6zm9). The five anti-parallel betta-strands do not form a barrel. *Lower panel:* Fold topology of Siv3-3 SAMase monomer. The lack of the second betta-pleated sheet that closes the barrel-like structure in T3 SAMase is apparent.

**Figure S5.**
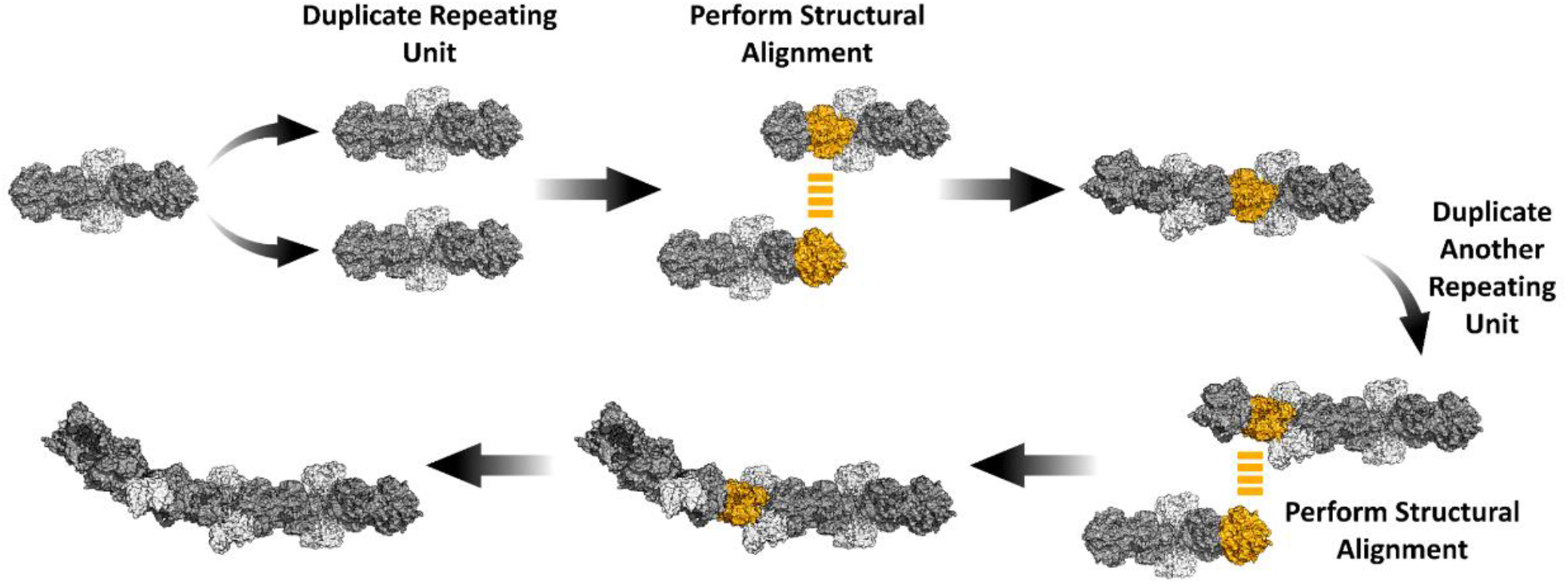
*In silico* reconstruction of SAMase-MAT filament. Filament reconstruction is done by iterative replication and alignment of SAMase-MAT hetero-oligomeric unit.

**Figure S6.**
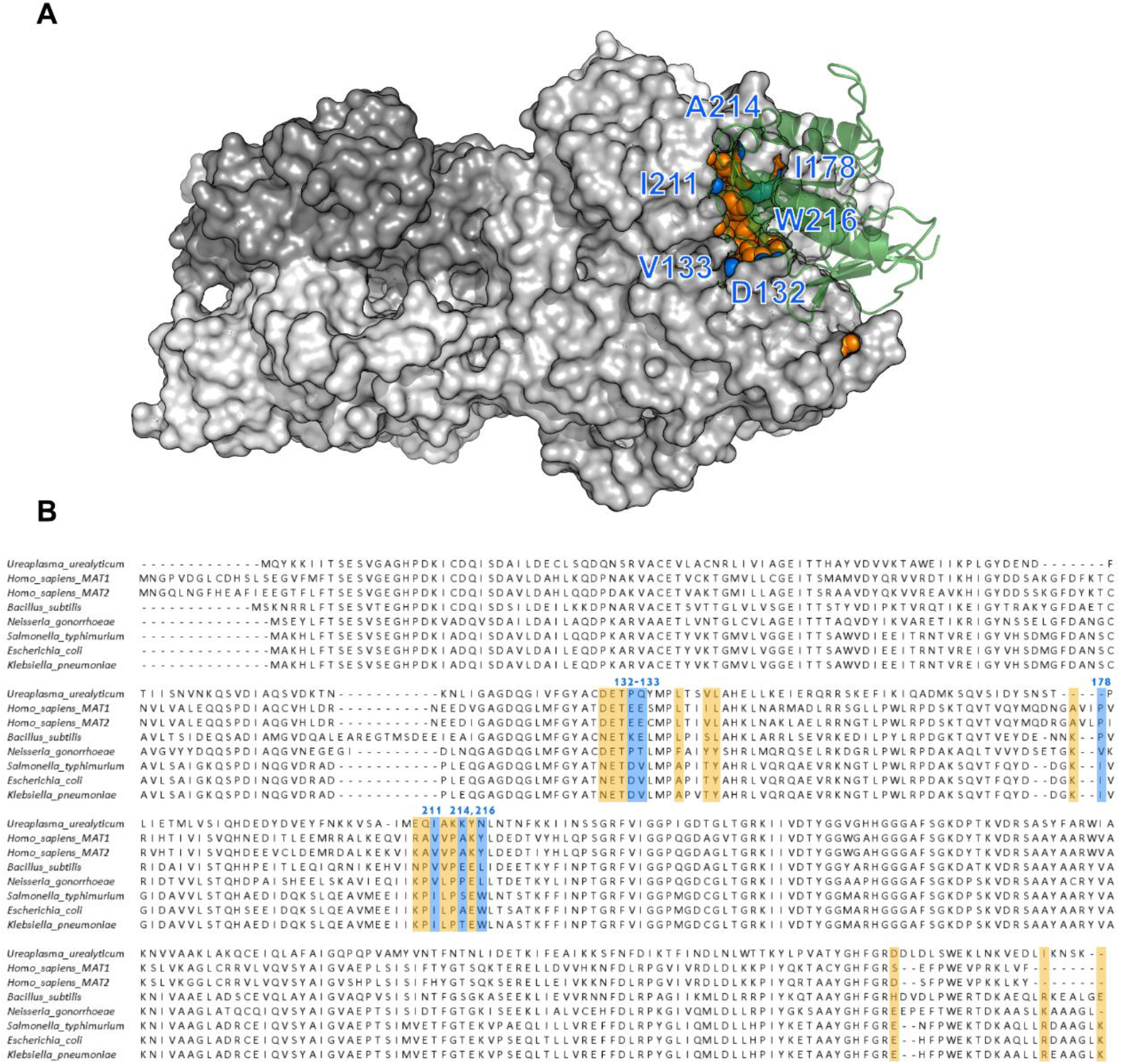
MAT residues forming interactions with SAMase. **A.** *E. coli* MAT residues forming an interface with a SAMase monomer are shown as *orange* and *blue* spheres. The coloring corresponds to the sequence alignment below. Residues in *blue* are found only in *E. coli* and other closely related enteric bacteria. SAMase monomer is shown in *green*. **B**. Sequence alignment of orthologous MATs. Residues involved in SAMase-*E. coli* MAT interface and their homologs in other MATs are colored in *orange* and *blue*. Residues replaced in *E. coli* MAT to break the interaction with SAMase are colored in *blue*. The residues are numbered according to *E. coli* MAT sequence.

**Figure S7.**
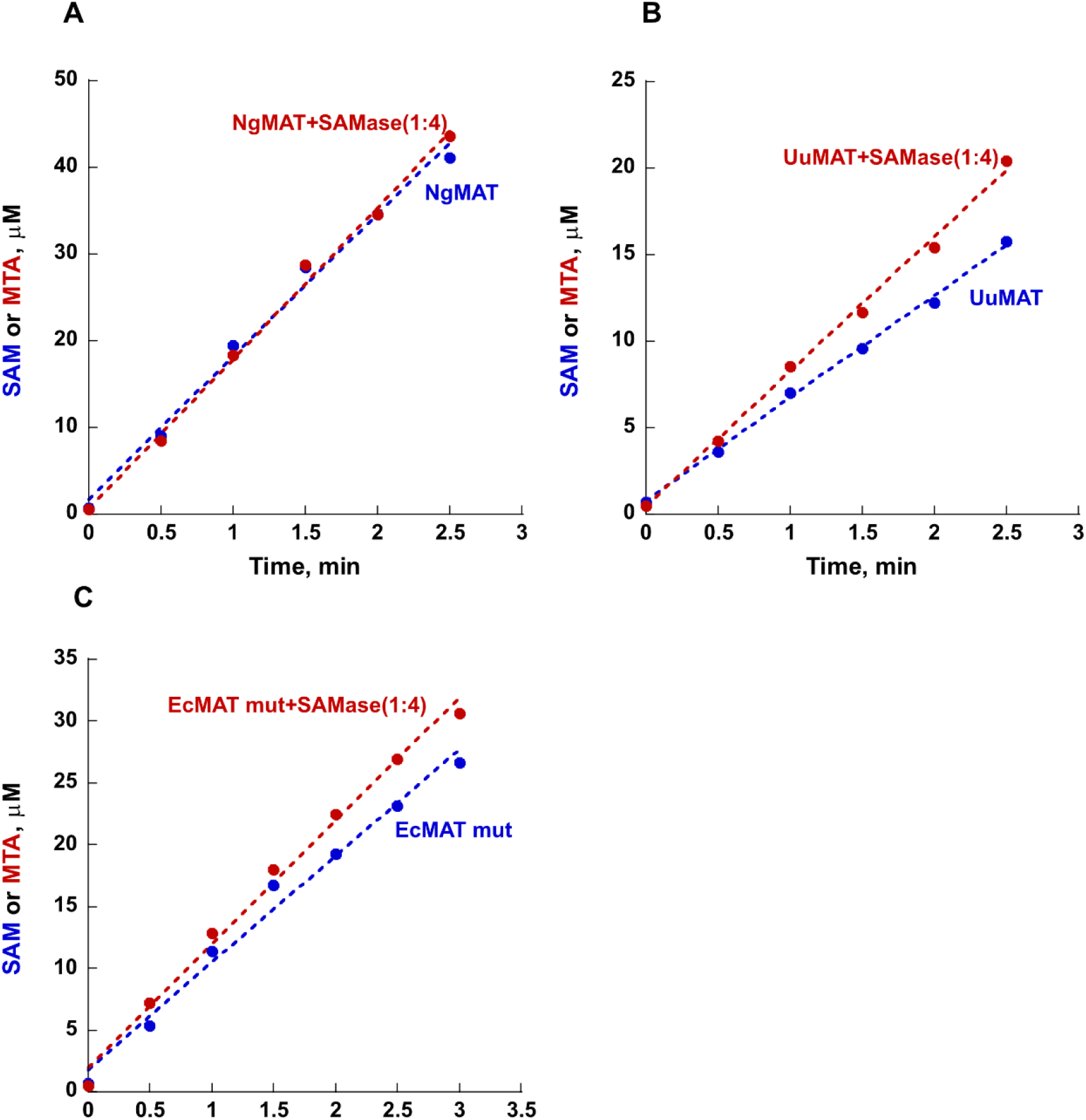
SAMase does not inhibit activity of distantly related orthologous MATs. Synthesis of SAM from ATP and methionine of MAT proteins was conducted in the presence of SAMase and the rate of SAM and/or MTA production was monitored by HPLC. **A, B.** Four-fold molar excess of SAMase relatively to *N. gonorrhea* MAT (NgMAT) (**A**) or *U. urealyticum* MAT (UuMAT) (**B**) did not inhibit MAT activity. **C.** Replacement of E. coli MAT (EcMAT) residues interacting with SAMase fully removes activity inhibition. EcMAT residues directly involved in SAMase-MAT interaction were replaced with their homologues in NgMAT (see **Fig. S6** for details), and the activity of the resulted mutant *E. coli* MAT (EcMAT mut) was measured either alone (*blue* trace, rate of SAM production from ATP and MET), or in the presence of four-fold molar excess of SAMase (*red* trace, MTA formation). The apparent increase in UuMAT and EcMAT mutant in the presence of SAMase is best explained by a reduction in product inhibition due to SAM degradation.

**Figure S8.**
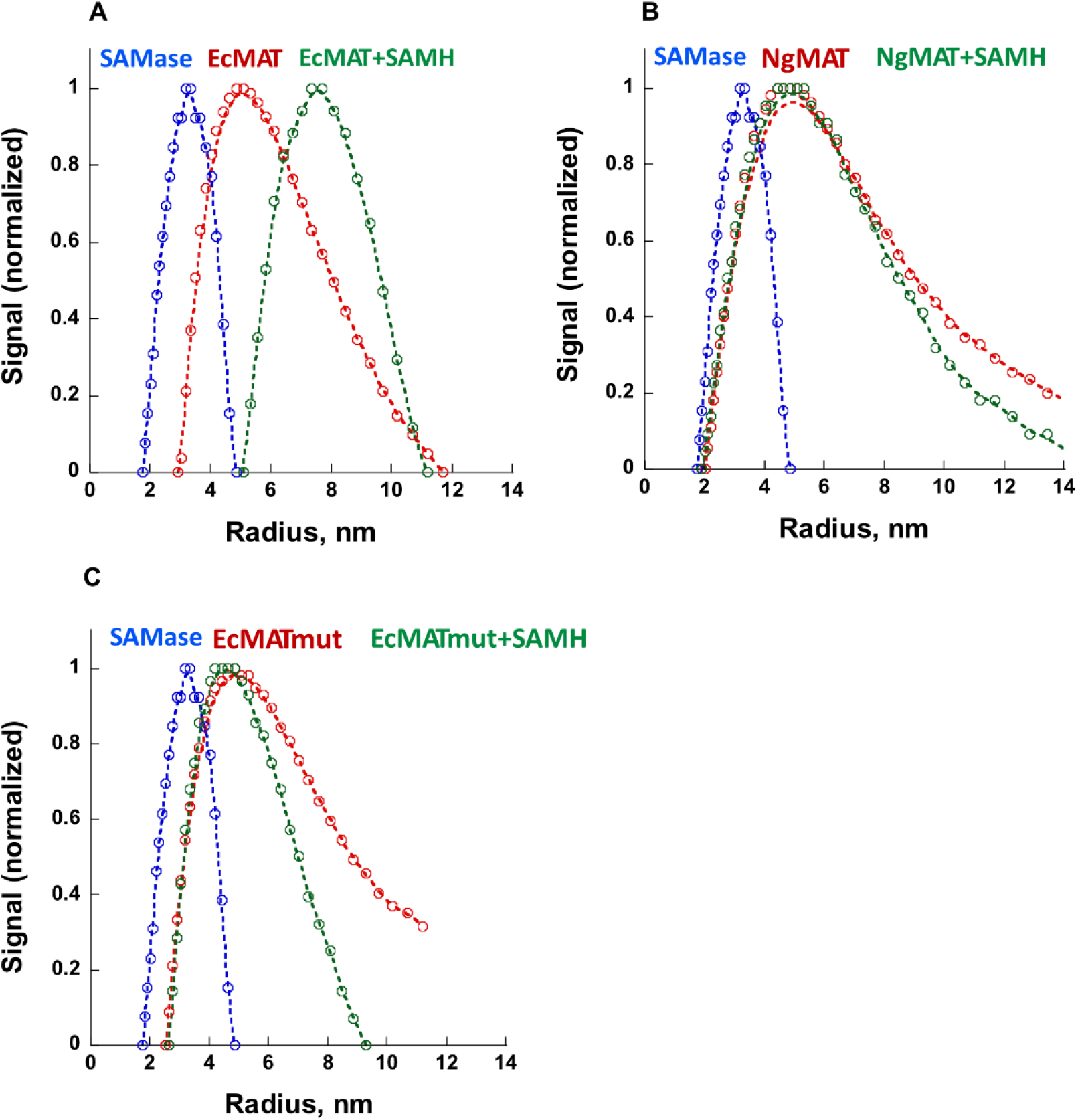
Dynamic Light Scattering (DLS) analysis of SAMase-MAT complex formation. SAMase, *E. coli* MAT (EcMAT), *E. coli* MAT to *N. gonorrhea* mutant (EcMAT mut), and *N. gonorrhea* MAT (NgMAT) were analyzed alone (*blue* and *red* traces) or as SAMase-MAT mix (*green* traces) after pre-incubation with SAM (see **Methods** for details). A clear shift in the calculated average radius is observed upon mixing of SAMase with EcMAT (**A**). No such shift can be seen in SAMase-NgMAT (**B**), or SAMase-EcMAT mut (**C**) mixes.

**Figure S9.**
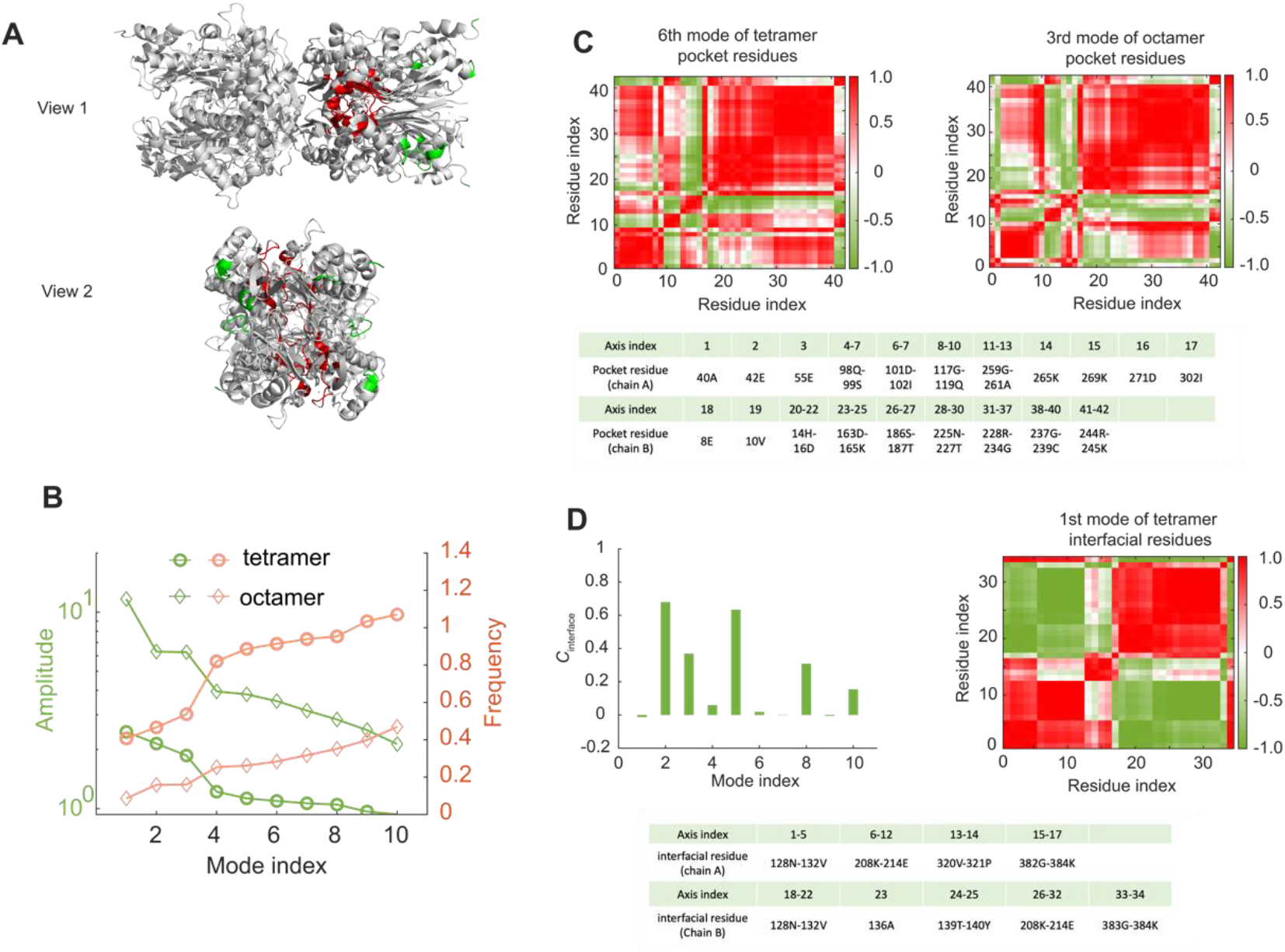
DMD simulations and normal mode analyses. **A.** Pocket residues (*red*) and interface residues (*green*). The conformation of tetramer is adopted from PDB databank (ID: 1P7L). **B.** Amplitude and frequency of motion of pocket (active site) residues as a function of mode index. ‘Tetramer’ corresponds to active site residues in an unbound MAT tetramer. ‘Octamer’ corresponds to active site residues within two MAT tetramers bound by two SAMase dimers. **C**. Correlation matrix of 6^th^ mode of tetramer pocket (*upper left panel*) and 3^rd^ mode of octamer pocket (*upper right panel*). Pocket residues are listed in the *lower panel*. Positive correlation means the pair-wise residues move in the same direction whereas negative correlation (anti-correlation) indicates that two residues move in the opposite direction, which leads to pocket deformation. **D.** *Upper left panel:* Mean inter-residue dynamic correlation of interface residues, *C_interface_*, as a function of normal mode index. Interface residues of tetramer undergo large motions at low-frequency modes. The low-frequency modes of tetramer involve large out-of-phase motions of the interface residues, which are inhibited or minimized upon polymerization. *Upper right panel:* Correlation matrix of 1^st^ mode of tetramer interface residues. Obvious anti-correlation can be found between interface residues of tetramer. Interface residues are listed in the *lower panel*.

**Figure S10.**
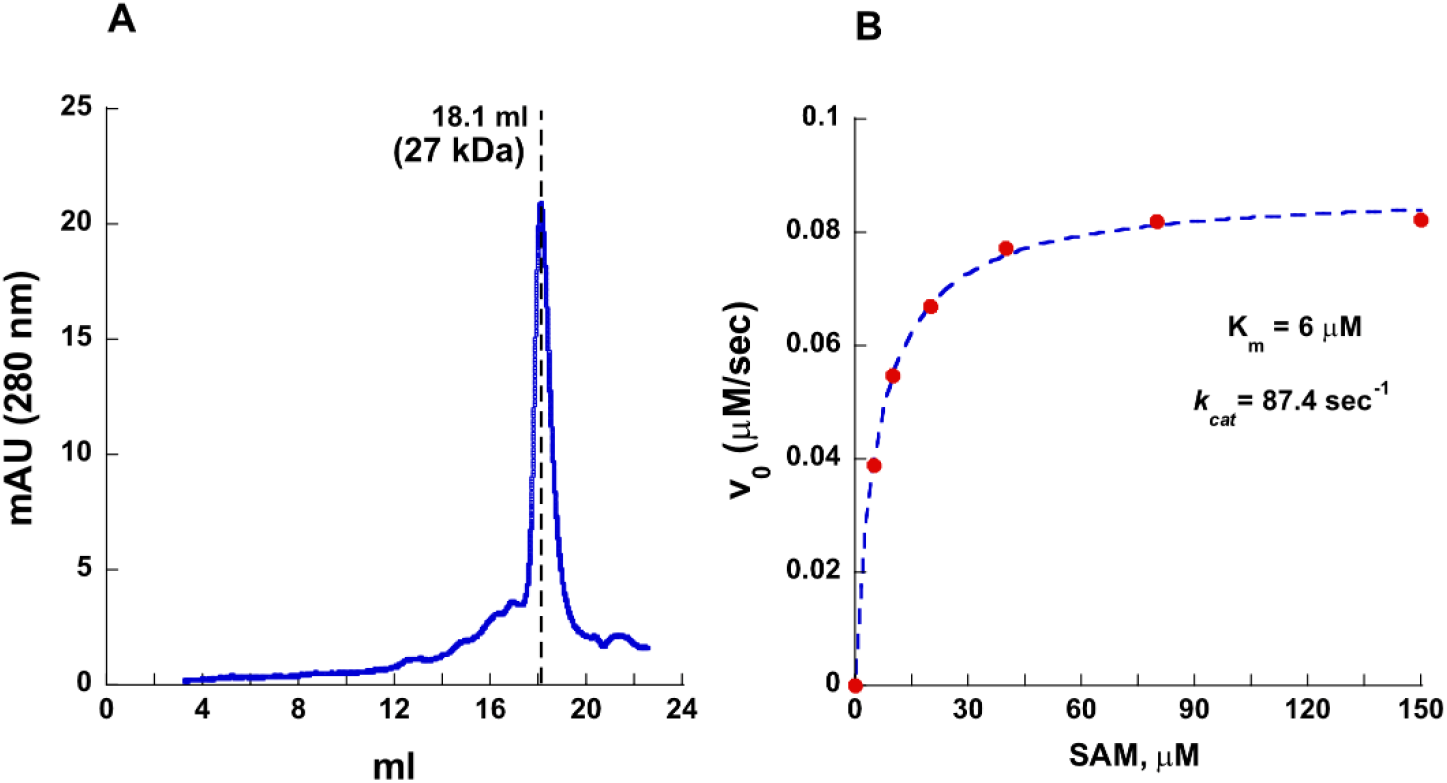
Characterization of refolded SAMase. **A**. Size Exclusion Chromatography of SAMase produces a single peak corresponding to a dimeric form. **B**. Michaelis-Menten fit of a refolded SAMase activity at a range of SAM concentrations.

**Figure S11.**
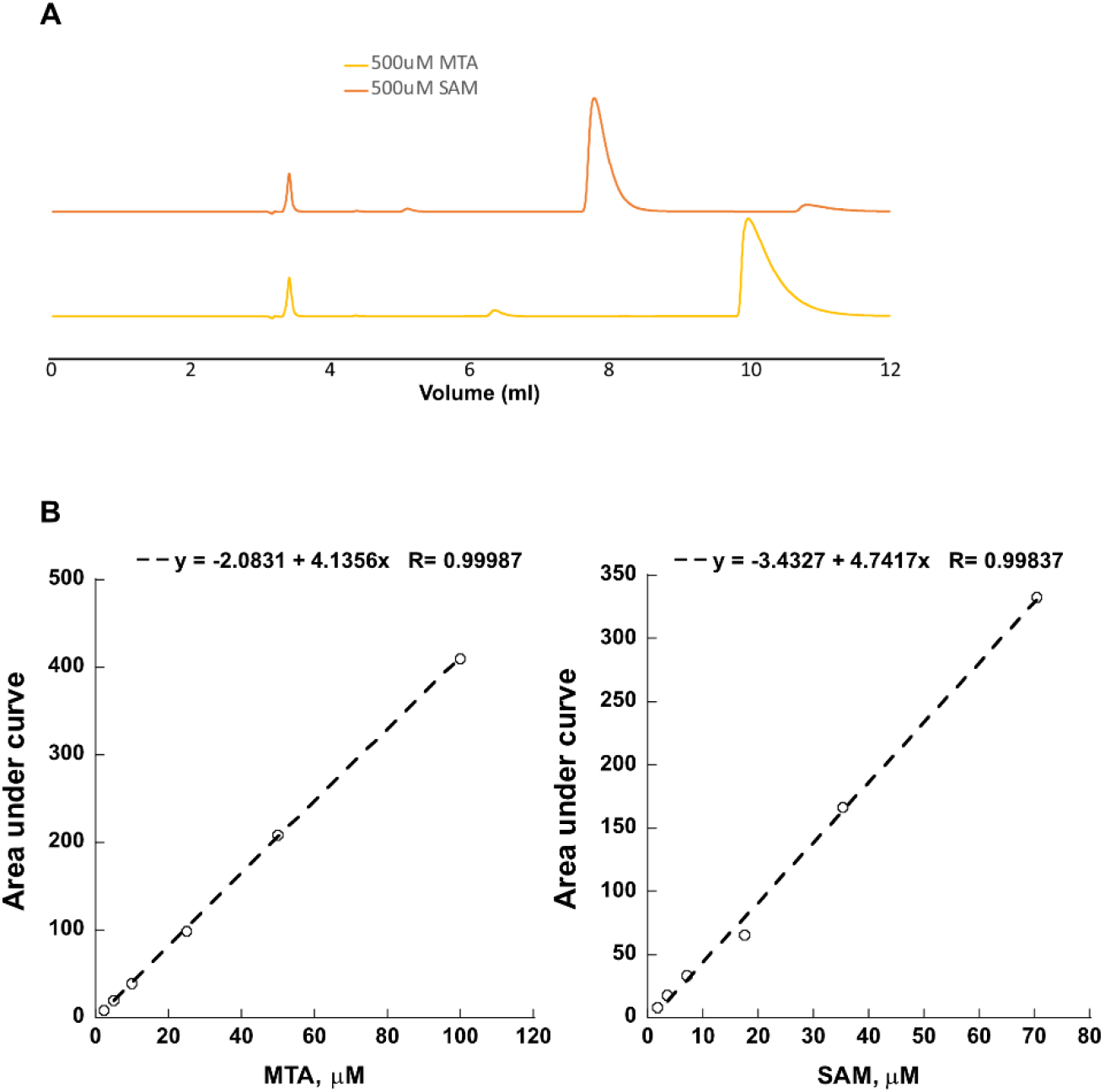
HPLC analysis. **A**. HPLC traces of 500μM SAM (*orange*) and 5’-methylthioadenosine (MTA) (*yellow*) standards. The standards were prepared in activity buffer and detected at 254 nm. **B.** The integrated area of SAM and MTA traces produced by HPLC separation strongly correlates with the injected concentration of standards.

